# PIWI proteins as prognostic markers in cancer: a systematic review and meta-analysis

**DOI:** 10.1101/663468

**Authors:** Alexios-Fotios A. Mentis, Efthimios Dardiotis, Athanassios G. Papavassiliou

**Affiliations:** Public Health Laboratories, Hellenic Pasteur Institute, Athens, Greece; Department of Microbiology, University Hospital of Thessaly, Larissa, Greece; Department of Neurology, University Hospital of Thessaly, Larissa, Greece; Department of Biological Chemistry, National and Kapodistrian University of Athens Medical School, Athens, Greece

**Keywords:** Cancer, Biomarkers, Prognosis, PIWI protein, human, survival analysis, bias, systematic review, meta-analysis, prognosis

## Abstract

**Background:** PIWI proteins, which interact with piRNAs, are implicated in stem cell and germ cell regulation, but have been detected in various cancers, as well.

**Objectives:** In this systematic review, we explored, for the first time in the literature (to our knowledge), the association between prognosis in patients with cancer and intratumoral expression of PIWI proteins.

**Data sources:** PubMed, Embase and Web of Knowledge databases were searched for the relevant cohort studies.

**Study eligibility criteria:** Prospective or retrospective cohort studies investigating the association of intratumoral mRNA or protein expression of different types of PIWI proteins with survival, metastasis or recurrence of various types of cancers in the systematic review. Exclusion of cross-sectional studies, of studies on the prognostic value of genetic polymorphism of PIWI genes, of studies re-analyzed previously published databases, and of conference abstracts and non-English articles.

**Participants:** Twenty-six studies with 4,299 participants were included in the systematic review.

**Interventions:** Pooled Hazard Ratios (HRs) and their 95% Confidence Intervals (CIs) were calculated for different PIWI proteins separately, by pooling of log of the calculated HRs using the random-effects model.

**Study appraisal and synthesis methods:** Data extraction was performed using a pre-designed form and quality of the studies was assessed using REMARK criteria. Heterogeneity assessed using the I2 index and the Cochran Q test. Publication bias assessed using funnel plots and Egger’s regression.

**Results:** The pooled HR of mortality in high compared to low expression of HIWI, HILI and PIWIL4 was 1.87 (CI95%: 1.31-2.66, p < 0.05), 1.09 (CI95%: 0.58-2.07, p = 0.79) and 0.44 (CI95%: 0.25-0.76, p < 0.05), respectively. The pooled HR of recurrence in in high compared to low expression of HIWI and HILI was 1.72 (CI95%: 1.20-2.49, p < 0.05) and 1.98 (CI95%: 0.65-5.98, p = 0.23), respectively.

**Limitations:** Exclusion of studies not in English; Discrepancy between mRNA and protein levels, and the respective analytical methods; Only one cancer site – PIWI protein pair investigated in three or more studies.

**Conclusions and Implications of Key Findings:** The prognosis of cancer patients is worse with higher HIWI and lower PIWIL4 expression, although the results are highly variable for different cancers. The expression of these proteins can be used for personalized prognostication and treatment of individual patients.

**Systematic review registration number:** Not registered.

## INTRODUCTION

Cancer prognosis using accurate, less clinical intuition-based approaches is pivotal during cancer patients’ management, and it forms the objective of several studies, including those applying machine learning approaches (Bertsimas et al., 2018) or the so-called “polygenic risk scores” (Mavaddat et al., 2019), which are based on SNPs in protein-coding genes. However, a tiny fraction of the human genome (<5%) is translated to proteins, while most of its regions are transcribed to non-coding RNAs (Ng et al., 2016), with potential implications in cancer.

Small regulatory non-coding RNAs, include microRNAs (miRNAs), small interfering RNAs (siRNAs) and P-element-induced wimpy testis (PIWI)-interacting RNAs (piRNAs). The piRNAs are non-coding RNAs that interact with Argonaute family proteins (PIWI proteins) to regulate gene expression. piRNAs are 25-31 nucleotides in length, i.e., a few nucleotides longer than miRNAs and siRNAs (Ng et al., 2016, Suzuki et al., 2012). PIWI proteins are a subclass of Argonaute proteins family that bind piRNAs to form piRNA-induced silencing complex (piRISC), which protects the genome’s integrity through silencing transposons (Han et al., 2017). Currently, four PIWI proteins have been identified in humans: PIWIL1 or HIWI, PIWIL2 or HILI, PIWIL3, and PIWIL4 or HIWI2 (Sasaki et al., 2003). PIWI proteins and piRNAs are normally expressed in the germline and are involved in spermatogenesis (Han et al., 2017).

Piwi-RNAs have been listed in the “dark matter of the genome”; however, their pathophysiological role is increasingly investigated in many fields, from cancer to neurodegeneration, and it is now known that the production of piRNAs is linked to the cleavage of the piRNAs-associated PIWI protein (Vourekas and Mourelatos, 2018) (Gainetdinov et al., 2018) (Sun et al., 2018) (Nandi et al., 2016). While previous studies (including systematic reviews and meta-analyses) have focused extensively on other RNA molecules, such as microRNAs (Nair et al., 2012, Wei et al., 2019, Cai et al., 2018), long non-coding RNAs (Serghiou et al., 2016) or circular RNAs (Li et al., 2018, Wang et al., 2018a, Chen et al., 2018, Wang et al., 2016) for either their diagnostic or prognostic role, the role of piRNAs in cancer is far less investigated, let alone in a systematic manner.

Nevertheless, two caveats exist before piRNAs potentially reach clinical prime as biomarkers: (i). Recent studies on cancer biomarkers have focused on both DNA and proteins to be used as biomarkers, highlighting therefore proteins’ role as biomarkers (Cohen et al., 2018) (Wang et al., 2018b). Using proteins as biomarkers may be equally suitable to, if not superior than, DNA or RNA, especially if proteins’ preservation during formalin overfixation at FFPEs is taken into account (Ammerlaan et al., 2018); and (ii). the number of piRNAs is estimated at ~20,000 in the eukaryotic genome; hence, this imposes a hurdle in easily identifying the most eligible among them as prognostic cancer biomarker.

On the other hand, PIWI proteins may be a more attractive biomarker: (i). there are four human paralogues of PIWI proteins in humans; (ii). these proteins are closely related to piRNAs pathophysiology; and (iii). they have been found to correlate with the characteristics of cancer stem cells (CSCs), including their potential for self-renewal and the ability to generate differentiated cells (Tan et al., 2015). In addition to the germline stem cells, where they are normally expressed, over-expression of PIWI proteins is noted in various human cancers, e.g., as in (Siddiqi and Matushansky, 2012, Stohr et al., 2019, Tosun et al., 2019). Furthermore, the PIWI proteins expression correlates with the tumor size, stage, differentiation grade, invasiveness as well as survival and recurrence (Eckstein et al., 2018).

However, no study enabling the systematic study and quantification of the prognostic role of PIWI proteins in cancer prognosis has been conducted so far, to our best knowledge. This prompted us to systematically review and meta-analyze here-in, for the first time, cohort studies that evaluated the association of PIWI proteins expression with mortality or recurrence of the tumor in patients with different types of cancer. The major limitation is that we combined all available data from all types of cancers in order to have sufficient number of studies included in the meta-analysis; this is a strategy that necessitates caution in the study’s conclusions. However, our study represents the first time in the literature to our best knowledge, that a meta-analysis in the field is conducted, and our overall goal was to investigate, at this stage, the role of PIWI proteins as prognostic marker in cancer, in general.

## METHODS

We performed this systematic review and meta-analysis according to the Preferred Reporting Items for Systematic Reviews and Meta-Analyses (PRISMA) statement (Moher et al., 2009, Liberati et al., 2009).

### Search strategy

PubMed, Embase and Web of Knowledge databases were searched for the relevant studies published from their inception until October 19^th^, 2018. EMBASE Classic was not searched necessarily because it contains studies published before 1973, i.e., before the first description of PIWI proteins and their homologues in non-human species (Lin and Spradling, 1997). Our search strategy was a combination of the terms referring to the PIWI proteins including PIWI, PIWIL, HIWI and HILI, and the terms related to cancer including itself, carcinoma, tumor, neoplasia, malignancy, metastasis, leukemia, lymphoma (Table S1), following previous suggestions for how to apply search string (e.g., (Mentis and Papavassiliou, 2018)) among others). We combined the results of the search in different databases and removed the duplicate records. No clinical queries prognosis filter, as previously described in (Haynes et al., 2005), was applied, i.e., all search results were curated by authors (AFAM and ED) reviewing them in total.

### Eligibility criteria

We included prospective or/and retrospective cohort studies that had investigated the association of intratumoral mRNA or protein expression of different types of PIWI proteins with survival, metastasis or recurrence of various types of cancers in the systematic review. We excluded cross-sectional studies that had compared PIWI expression in cancerous and normal tissue with no data on survival, studies that had explored the prognostic value of genetic polymorphism of PIWI genes (such as (Sung et al., 2012, Zhang et al., 2016)), studies that had analyzed a previously published database (such as (Krishnan et al., 2016, Xie et al., 2018)), conference abstracts and non-English articles. We also compared the list of authors, centers of study, and recruitment intervals of the included studies to identify the duplicate reports of the same research or studies with a substantial overlap in their study population. For such cases, the less informative studies were excluded.

### Study selection

To identify the relevant studies, we first screened the titles and/or abstracts of the combined search results and selected the certainly or possibly relevant studies for the second stage of the study selection process. In the second stage, after excluding the conference-abstracts, we retrieved the full-texts of these studies, evaluated them in detail and included the studies which satisfactorily met the eligibility criteria. To eliminate personal bias, the whole search strategy process was conducted independently by two reviewers (AFAM and ED) who evaluated the title, abstract as well as full text of the papers on the basis of the above defined inclusion and exclusion criteria; any disagreements were discussed afterwards with a third author (AGP).

### Data extraction and quality assessment

We designed a data extraction spreadsheet on Google Sheets (with input from the CHARMS checklist (Moons et al., 2014)) and for each included study recorded the following data fields: first author, country, study type, number of participants, cancer site and type, important inclusion and exclusion criteria, age, gender, clinical stage, differentiation grade and TNM of the tumors, treatment details, follow-up duration and protocol, mortality outcomes definition and rate including overall survival (OS) and disease-specific survival (DSS), recurrence or metastasis outcomes definition and rate including recurrence free survival (RFS), progression or disease free survival (PFS/DFS) and metastasis free survival (MFS), method of PIWI expression measurement, and the statistical analyses.

We used the Reporting Recommendations for Tumor Marker Prognostic Studies (REMARK) checklist to assess the quality of the included studies (Altman et al., 2012) and its updated, abridged version (Sauerbrei et al., 2018); this checklist is part of the EQUATOR Network guidelines (Simera et al., 2010). For individual studies, each of the 20 items of this checklist were marked as fully satisfied, partially satisfied, not satisfied, unclear, or not applicable. The fully satisfied items were scored with one and partially satisfied items with half points, and the quality of each study was scored as percentage by dividing the sum of scores for each item by the number of applicable items. We did not exclude any study based on its quality score to avoid bias.

### Statistical analysis

Studies were included in the meta-analysis if they had reported RR, HR or Kaplan-Meier curve. For the eligible studies that univariable HRs were not reported, we extracted the data points of the Kaplan-Meier curves using Plot Digitizer 2.6.8, and we estimated the univariable HR, following previous publications (Tierney et al., 2007a). Following the example of previous studies (Serghiou et al., 2016), if three or more studies were available for each analysis, we performed separate meta-analyses for different types of PIWI proteins comparing risk of outcomes in high compared to low as well as intermediate compared to high or low PIWI expression regardless of the cancer types. When available, multivariable HR or RR, and otherwise, univariable HR or RR was selected as the effect size for each study.

The meta-analyses were performed using Comprehensive Meta-Analysis 2.2 (BioStat Inc., US). The pooled HRs, on the basis of the Mantel-Haenszel method, and their 95% CIs were calculated using the random-effects model, as in (Tierney et al., 2007b). We considered a meta-analysis heterogeneous when the I^2^ index was greater than 70% and the Cochran Q test was statistically significant (Higgins et al., 2003). Subgroup analyses were initially planned to be performed based on cancer sites, method of PIWI expression measurement, outcome type, method of analysis (univariable or multivariable), and whether detectable and undetectable PIWI expressions are compared, on the condition that three or more studies were available for each subgroup. Publication bias, which reflects the tendency of publishers and authors to publish statistically significant results, was examined using funnel plots and Egger’s regression (Egger et al., 1997). To create the funnel plots, the standard error was plotted against the natural logarithm of hazards ratio. To assess this bias, if more than 10 studies were included in a meta-analysis, we visually inspected its funnel plot and performed Egger’s regression; asymmetric funnel plot and statistically significant Egger’s regression represented the publication bias. Statistical significance was set at p<0.05.

### Re-analysis of data from the *Pathology Atlas of The Human Cancer Transcriptome*

To investigate the pattern of expression of PIWI-like proteins across various cancer types and their effect on patient survival, we mined the Human Protein Pathology Atlas (https://www.proteinatlas.org/humanproteome/pathology) (Uhlen et al., 2017). For each of the PIWIL proteins corresponding genes, normalized expression values (log2 FPKM) from the TCGA (Uhlen et al., 2017) are presented as box plots covering 17 broad cancer types.

## RESULTS

Our search collectively revealed 1,538 records. After removing overlaps between the databases, the titles and/or abstracts of the remaining 826 records were screened. We identified 90 relevant records from which 21 were abstracts and the remaining 69 manuscripts were examined in detail. Ultimately, 26 studies with 4,299 participants were included in the systematic review (Al-Janabi et al., 2014, Cao et al., 2016, Chen et al., 2015, Gambichler et al., 2017, Greither et al., 2012, Grochola et al., 2008, He et al., 2009, Iliev et al., 2016, Li et al., 2012, Li et al., 2017, Litwin et al., 2018, Liu et al., 2012, Lu et al., 2016, Navarro et al., 2015, Oh et al., 2012, Pouyanfar et al., 2016, Qu et al., 2015, Sun et al., 2011, Sun et al., 2017, Taubert et al., 2007, Taubert et al., 2015, Wang et al., 2012, Zeng et al., 2017, Zeng et al., 2011, Zhang et al., 2013, Zhao et al., 2012) (Figure 1, Table 1).

**Figure 1.**
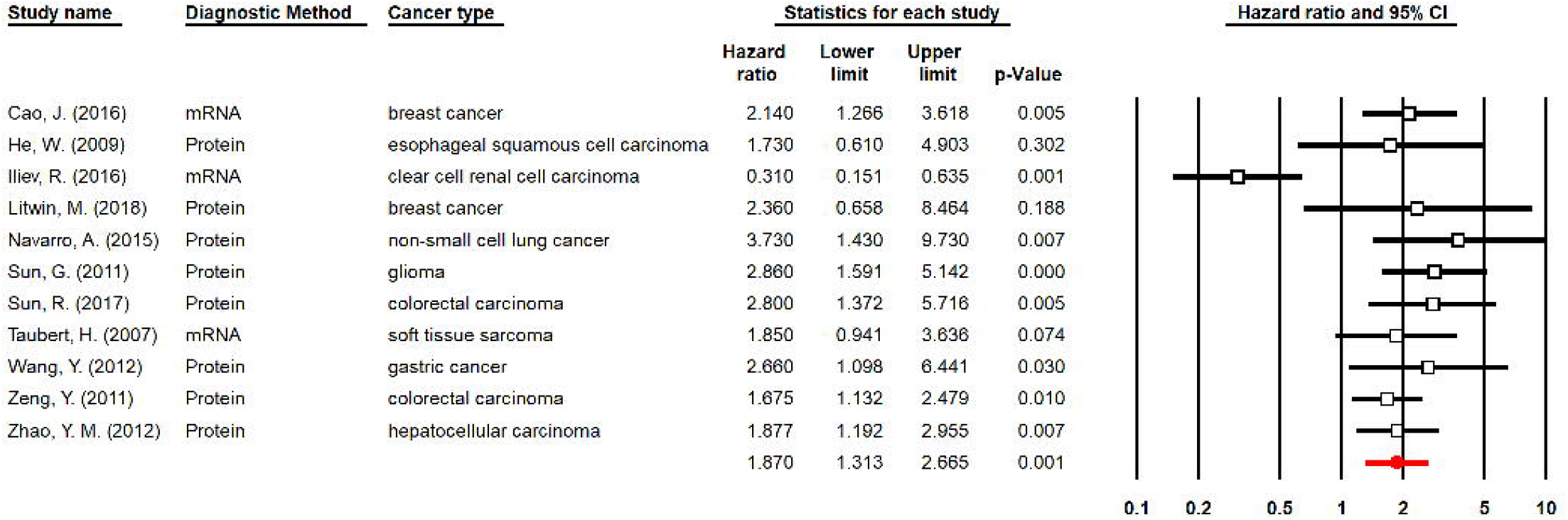
The study selection process. The searched revealed 1,538 records. After removing 712 overlaps between the databases, the remaining 826 records were screened and 90 relevant records were identified, from which 21 were conference abstracts. The remaining 69 manuscripts were examined using their full-text and 26 eligible studies were identified.

The HIWI (PIWI1), HILI (PIWI2), PIWI3, and PIWI4 protein were studied in 15, 15, 5, and studies, respectively. Colorectal cancer with five studies, and breast cancer with three studies were the most studied types of cancer. The included studies were performed in different countries in Asia and Europe, mostly in China and Germany. Twenty-five studies were cohorts (including four retrospective, one prospective and 20 unspecified), and one was a case-control study. The average age of the participants was in the range of 48 to 68 years across the studies and was not reported in three studies. The ratio of female patients in the cancers which are not sex-specific was widely variable from 6.5% to 54.6%. Early and advanced cases of cancer were evenly included in the systematic review, with 29% in the stages I and II and 25% in the stages III and IV, while the clinical stage was not reported for 45.1% of the patients. The rates of lymph node and distant metastasis reported in seven and three studies, was 42% and 13%, respectively. While the majority of the patients did not receive neoadjuvant treatments, four studies reported using adjuvant chemotherapy. The expression of PIWI genes was measured using immunohistochemistry (IHC) in 18 studies and reverse transcription PCR (RT-PCR) in eight studies. Patients were followed on average from 16 to 124 months, although the duration of follow-up was not reported in nine studies. OS was investigated in 13, DSS in six, both OS and DSS in one, unspecified mortality in five, DFS or PFS in seven, RFS in two and unspecified recurrence in two studies (Table 2).

### HIWI (PIWIL1)

Fifteen studies including 1,862 patients investigated the prognostic value of HIWI expression in different types of cancers. The outcome of interest was mortality in 16 studies, and recurrence in five studies. Six studies on breast cancer, gastric cancer, glioma, esophageal squamous cell carcinoma, colorectal cancer, hepatocellular carcinoma and non-small cell lung carcinoma reported significantly higher risk of mortality with higher HIWI expression. In multivariable analyses, four studies showed that HIWI remains a significant predictor of increased mortality after considering confounding variables such as tumor size, clinical stage, differentiation grade and PIWI2 expression. Conversely, one study on clear cell renal carcinoma showed 70% lower risk of death in patients with higher HIWI expression. No significant difference in mortality between high and low HIWI expression was reported in five studies investigating patients with breast cancer, epithelial ovarian cancer, clear cell renal carcinoma, ductal adenocarcinoma of the pancreas and soft tissue sarcomas.

Eleven studies including 1,331 patients were eligible for the meta-analysis of mortality HR comparing high and low HIWI expression (Figure 2). The pooled HR was calculated as 1.87 (CI95%: 1.31-2.66, p < 0.05) while a moderate to high heterogeneity was observed between the studies (I^2^ = 67%, p < 0.05). The reported HRs were significantly different for various cancer sites. Subgroup analysis showed that the pooled HR for studies measuring mRNA and protein expression of HIWI was 1.09 (CI95%: 0.34-3.45, p = 0.87) and 2.12 (CI95%: 1.70-2.65), respectively. Additional subgroup analyses based on the outcome type, method of analysis, and whether HR was calculated or reported, showed no significant difference between the subgroups. Publication bias was not evident in this meta-analysis (Figure 3, p = 0.82).

**Figure 2.**
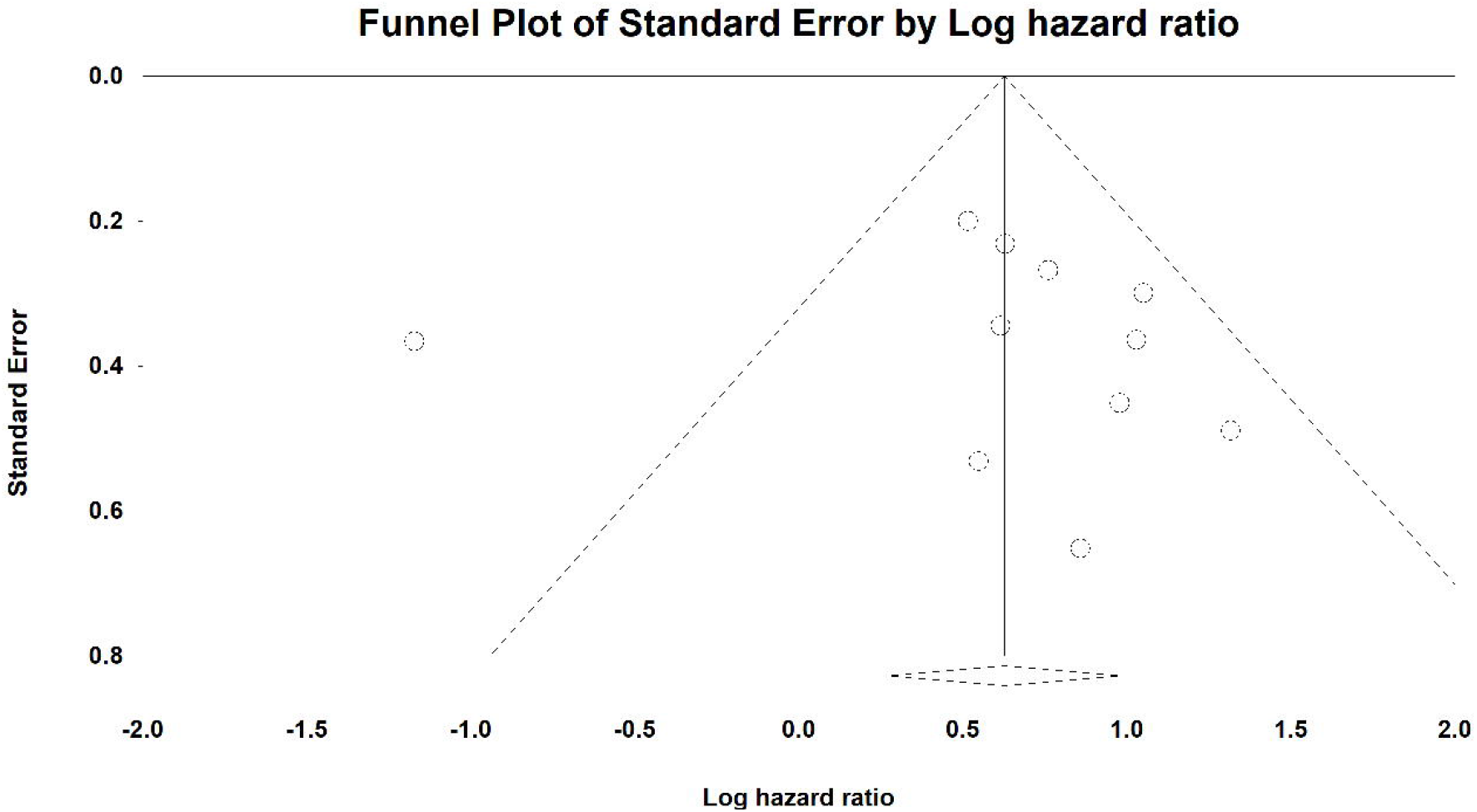
Meta-analysis of HR for mortality in high compared to low expression of HIWI (1,331 patients). Prognostic value of HIWI for mortality in individual studies are shown in each row. The hazard ratios for individual studies are denoted by squares bounded by a 95% confidence interval bar. The pooled hazard ratio is denoted by a red diamond bounded by a 95% confidence interval bar. Higher HR represents higher risk of mortality in patients with high HIWI expression.

**Figure 3.**
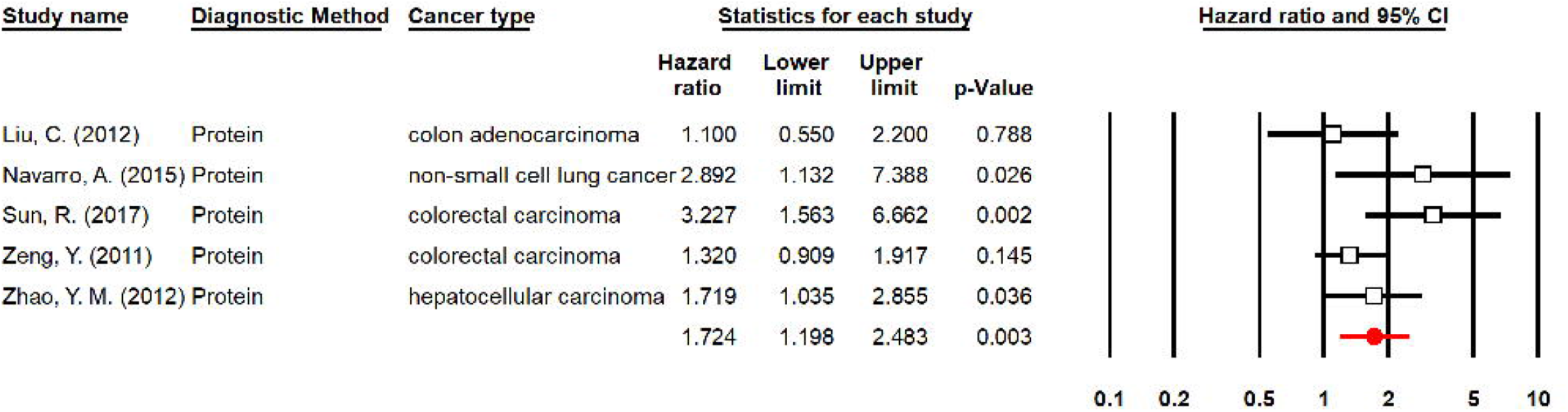
Funnel plot for the meta-analysis of HR for mortality in high compared to low expression of HIWI. The standard error is plotted against the natural logarithm of hazards ratio. Each study is represented by a circle and the vertical line shows the natural logarithm of pooled hazards ratio. Publication bias was not observed as shown by the symmetry of the plot and the Egger’s regression asymmetry test (p = 0.82).

The patients were classified into low, intermediate and high HIWI expression groups in four studies. Two studies on ductal adenocarcinoma of the pancreas and epithelial ovarian cancer showed a higher risk of mortality in patients with intermediate HIWI expression compared to its high or low expression (Grochola et al., 2008, Lu et al., 2016). Similarly, the estimated HR for another study showed a higher risk of mortality for intermediate HIWI expression in esophageal squamous cell carcinoma. Taubert et al. studied patients with soft tissue sarcomas and showed a significantly lower risk mortality in intermediate HIWI expression compared to its high but not low expression (Taubert et al., 2007). Three studies (429 patients) were included in the meta-analysis of mortality HR comparing intermediate HIWI expression with high or low HIWI expression. The pooled HR was 1.21 (CI 95%: 0.58-2.52, p = 0.607) while the meta-analysis was highly heterogeneous (I^2^ = 84%, p < 0.05).

Although the combined cytoplasmic and nuclear HIWI expression was investigated in most studies, one study evaluated the prognostic value of cytoplasmic and nuclear HIWI expression separately. They reported a significantly higher risk of mortality for cytoplasmic but not nuclear HIWI expression in patients with esophageal squamous cell carcinoma (He et al., 2009). In addition to intratumoral expression of HIWI, two studies measured its expression in the adjacent non-cancerous tissue as well. One study on colorectal carcinoma showed a significantly lower survival in patients with higher peritumoral HIWI expression (Zeng et al., 2011), but the other study on hepatocellular carcinoma showed no significant association (Zhao et al., 2012).

Six studies investigating 1,055 patients with ovarian, colorectal, hepatocellular and non-small cell lung cancers evaluated the association of recurrence with HIWI expression. Three studies showed a significantly higher recurrence in patients with higher HIWI expression after considering confounding variables such as clinical stage, differentiation grade and PIWI4 expression. Conversely, three studies on patients with colorectal and ovarian cancers reported no significant association between recurrence and HIWI expression. The pooled HR of recurrence in high compared to low HIWI expression was 1.72 (CI95%: 1.20-2.49, p < 0.05, Figure 4). Minimal heterogeneity was observed in this meta-analysis (I^2^ = 45%, p = 0.12). The pooled HR for the subgroup of three studies investigating colorectal cancer was 1.61 (CI95%: 0.92-2.82, p = 0.10). One study classified the patients into low, intermediate and high HIWI expression group and showed no significant difference in recurrence risk between intermediate and high or low HIWI expression (Lu et al., 2016).

**Figure 4.**
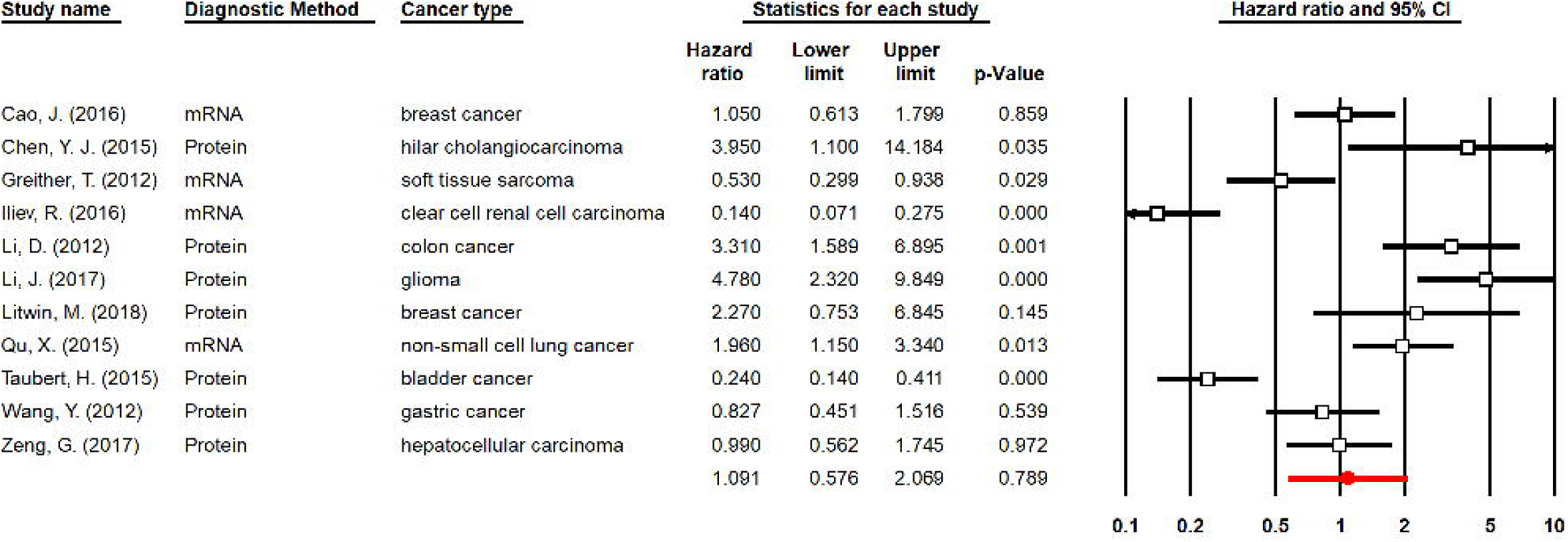
Meta-analysis of HR for recurrence in high compared to low expression of HIWI (844 patients). Prognostic value of HIWI for recurrence in individual studies are shown in each row. The hazard ratios for individual studies are denoted by squares bounded by a 95% confidence interval bar. The pooled hazard ratio is denoted by a red diamond bounded by a 95% confidence interval bar. Higher HR represents higher risk of recurrence in patients with high HIWI expression.

As shown in the re-analysis of the TCGA Atlas, PIWIL1 showed the largest range of expression with a relatively large number of samples across cancer types exhibiting expression above 20 FPKM. In line with the expression pattern of PIWIL1 in normal tissues, testicular cancer had the largest proportion of samples with high PIWIL1 expression. While most cancer types displayed very low expression of PIWIL1 (i.e., very few samples with expression above 5 FPKM), its expression was elevated in a substantial number of samples in thyroid, stomach and colorectal cancers, along with testicular cancer (Supplementary Figure 1).

### HILI (PIWIL2)

The prognostic value of HILI expression was evaluated in 15 studies including 2643 patients. Mortality was investigated in 15 and recurrence in five studies.

High HILI expression was associated with significantly lower survival in six studies investigating colorectal cancer, non-small cell lung cancer, gastric cancer, hilar cholangiocarcinoma, glioma, and prostate cancer. Additionally, another study on colorectal cancer showed a borderline non-significant association of HILI with mortality. HILI remained a significant predictor of mortality after considering confounding variables in two studies; however, in one study it failed to remain significant after including HIWI expression in the model (Wang et al., 2012). On the other hand, three studies on clear cell renal carcinoma, soft tissue sarcomas and bladder cancer showed a better survival in patients with higher HILI expression. In addition to the combination of cytoplasmic and nuclear HILI expression, one study investigated their prognostic effect separately and showed that the cytoplasmic but not nuclear HILI expression was associated with mortality (Taubert et al., 2015).

We included eleven studies with 1,389 participants in the meta-analysis of mortality HR in high HILI expression compared to low HILI expression (Figure 5). The pooled HR was 1.09 (CI 95%: 0.58-2.07, p = 0.79) in a substantially heterogeneous meta-analysis (I^2^ = 91%, p < 0.05). After excluding two studies that compared detectable and undetectable HILI expression (rather than high and low HILI expression), the pooled HR was 1.32 (CI95%: 0.65-2.70, p = 0.44). Subgroup analysis showed that the estimated HR is significantly different in various cancer sites and for different types of outcome. The pooled HR for DSS was 0.35 (CI95%: 0.16-0.77, p < 0.05) while it was 1.51 (CI95%: 0.53-4.35, p = 0.44) for OS. The pooled HR for studies measuring mRNA and protein expression of HILI was 0.63 (CI95%: 0.22-1.79, p = 0.39, p = 0.39) and 1.53 (CI95%: 0.64-3.67, p = 0.34), respectively. The effect of publication bias was not shown in this meta-analysis (Figure 6, p = 0.25).

**Figure 5.**
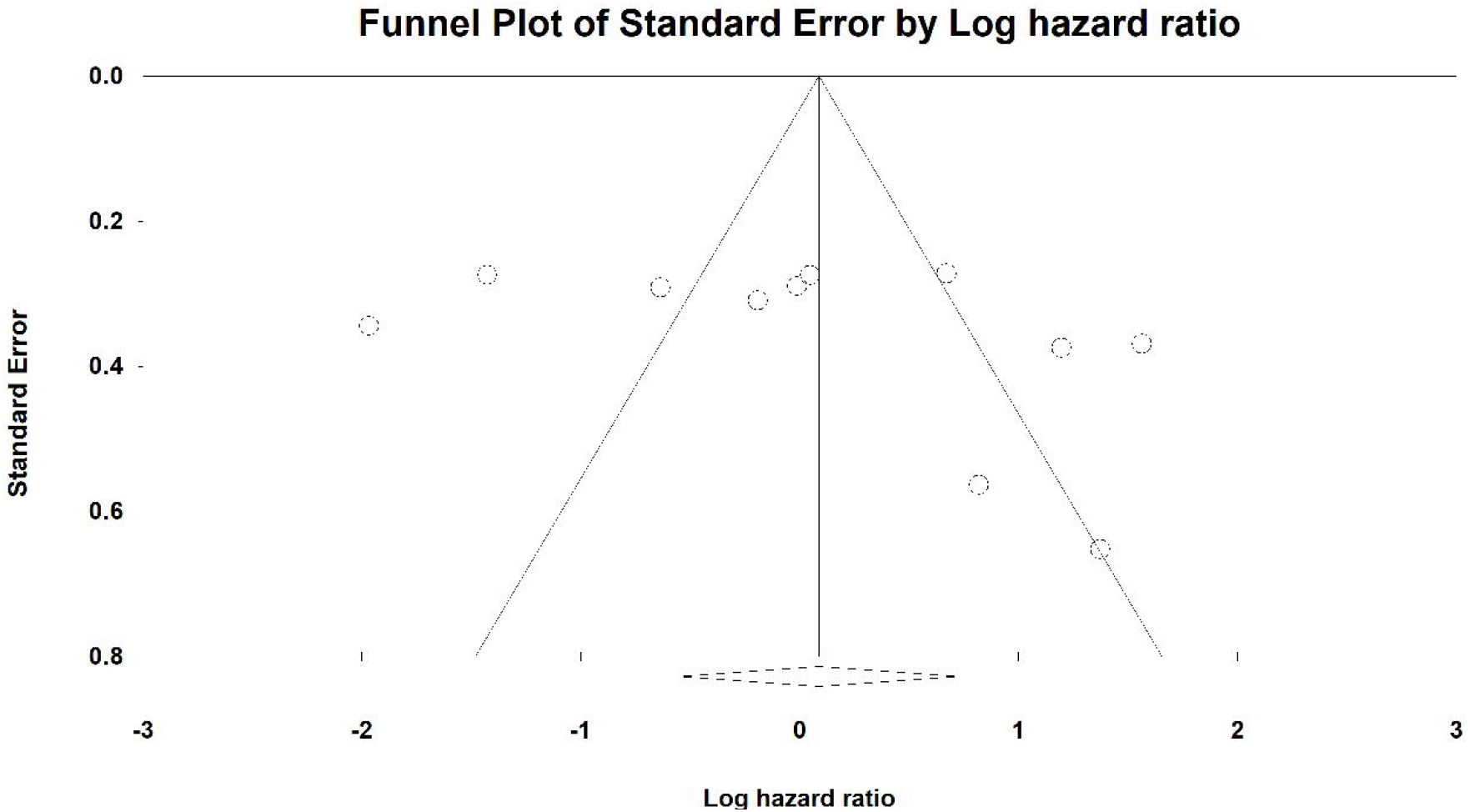
Meta-analysis of HR for mortality in high compared to low expression of HILI (1,389 patients). Prognostic value of HILI for mortality in individual studies are shown in each row. The hazard ratios for individual studies are denoted by squares bounded by a 95% confidence interval bar. The pooled hazard ratio is denoted by a red diamond bounded by a 95% confidence interval bar. Higher HR represents higher risk of mortality in patients with high HILI expression.

**Figure 6.**
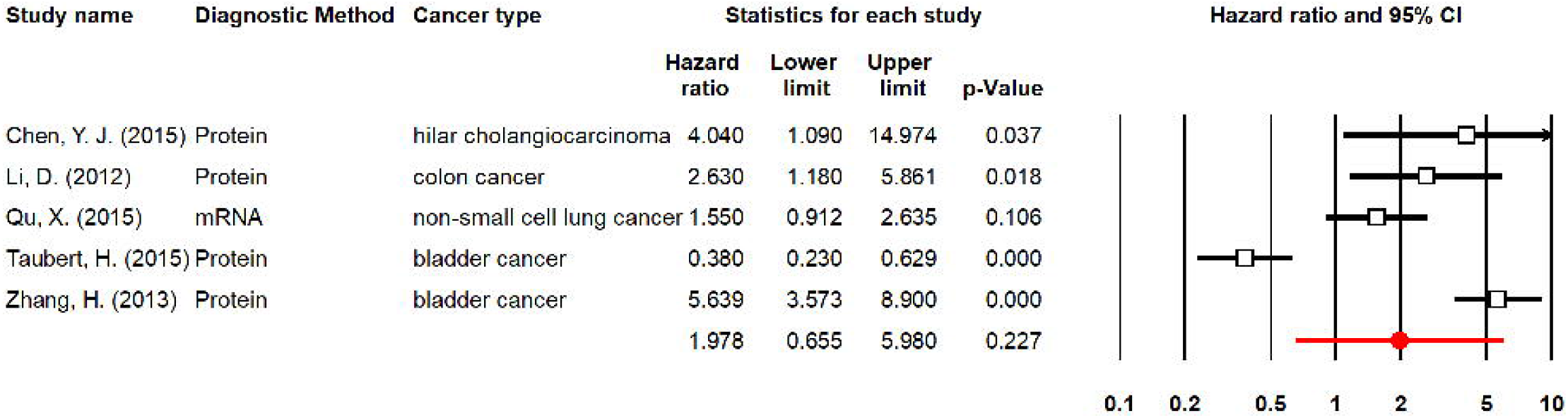
Funnel plot for the meta-analysis of HR for mortality in high compared to low expression of HILI (1,640 patients). The standard error is plotted against the natural logarithm of hazards ratio. Each study is represented by a circle and the vertical line shows the natural logarithm of pooled hazards ratio. Publication bias was not observed as shown by the symmetry of the plot and the Egger’s regression asymmetry test (p = 0.25).

While four studies showed a significantly higher risk of local or distant recurrence with higher HILI expression in patients with colorectal, breast, bladder and non-small cell lung cancers, one study on colorectal cancer showed no association between HILI expression and the risk of recurrence. In contrast, another study on bladder cancer showed that the patients with detectable HILI expression are at a lower risk of recurrence. They also investigated the nuclear and cytoplasmic HILI expression separately and showed the same effect in each component (Taubert et al., 2015). We performed a meta-analysis on five studies (including 1,640 patients) to calculate the HR of recurrence in high compared to low HILI expression (Figure 7). The pooled HR was 1.98 (CI95%: 0.65-5.98, p = 0.23) and the meta-analysis was quite heterogeneous (I^2^ = 94%, p < 0.05). Sensitivity analysis showed that after excluding the one study that compared detectable and undetectable HILI expression, the HR was 3.05 (CI 95%: 1.47-6.30, p < 0.05). The HRs for various cancer sites was significantly different. However, the pooled HR of 4.15 (CI95%: 1.99-8.63) for MFS was not significantly different from the pooled HR of 1.20 (CI95%: 0.35-4.18) for DFS. Additionally, the pooled HR of four studies which used protein expression of HILI was not significantly different from the one study which used its mRNA expression. Publication bias was not assessed due to the small number of included studies in this meta-analysis.

**Figure 7.**
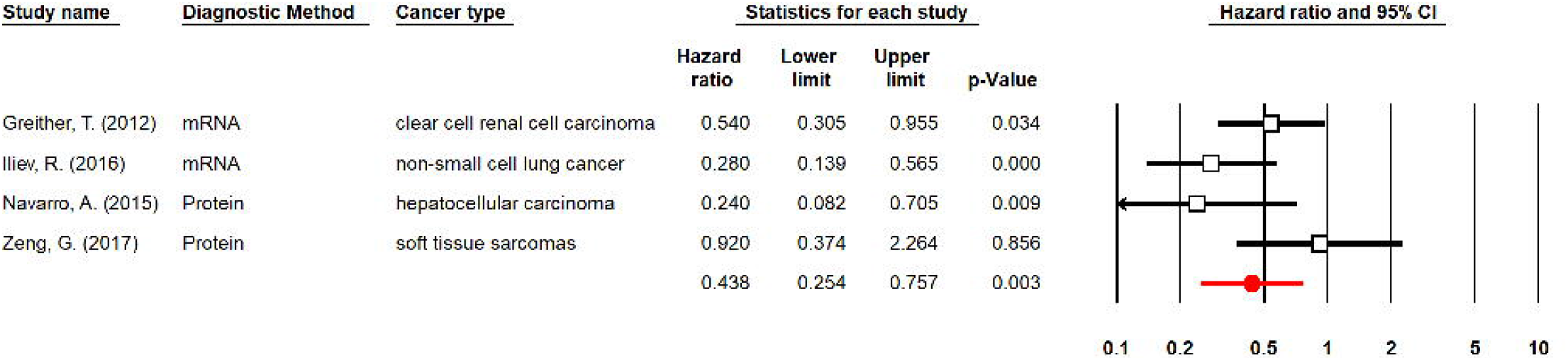
Meta-analysis of HR for recurrence in high compared to low expression of HILI. Prognostic value of HILI for recurrence in individual studies are shown in each row. The hazard ratios for individual studies are denoted by squares bounded by a 95% confidence interval bar. The pooled hazard ratio is denoted by a red diamond bounded by a 95% confidence interval bar. Higher HR represents higher risk of recurrence in patients with high HILI expression.

Based on the TCGA atlas, testicular cancer was by far the highest expressing cancer type for PIWIL2 gene. Other cancers with elevated PIWIL2 expression included Testis, Stomach and Colorectal. We also noted a slightly increased expression in Liver, Pancreatic and Head and Neck cancers. While displaying overall low expression, lung and breast cancers contained a large number of “outlier” samples with increased PIWIL2 expression (Supplementary Figure 2).

### PIWIL3

The association of PIWIL3 expression with the prognosis of clear cell renal carcinoma, malignant melanoma, soft tissue sarcomas and gastric cancer was investigated in five studies including 619 patients. All five studies that investigated the association of mortality with PIWIL3 expression found no significant association. While the statistics and method of analysis was not clear in two studies, only one study performed Cox regression and a meta-analysis was not performed. Greither et al. showed that the multivariable HR of mortality in high compared to low PIWIL3 expression was 0.75 (CI95%: 0.43-1.33, p = 0.33) (Greither et al., 2012). Only one study evaluated the association of recurrence with PIWIL3 expression on malignant melanoma and showed no significant association, while they failed to report the statistical analysis they used to investigate this association (Gambichler et al., 2017).

In the TCGA Atlas, PIWIL3 exhibited the lowest expression levels of the four PIWIL genes, with very few samples exceeding FPKM of 5 and majority of samples not exceeding FPKM of 1. Overall higher expression of PIWIL3 was noted in testicular cancer and an elevated number of “outlier” samples with higher PIWIL3 expression in lung and urothelial cancer (Supplementary Figure 3).

### PIWIL4

Six studies with 594 participants investigated the prognostic value of PIWIL4 in clear cell renal carcinoma, hepatocellular carcinoma, soft tissue sarcomas, gastric cancer, and non-small cell lung cancer. Mortality was studied in six studies and recurrence in one study. In three studies, significantly lower risk of mortality was observed in patients with higher PIWIL4 expression. These studies investigated patients with cell renal carcinoma, soft tissue sarcomas, and non-small cell lung cancer. One study on non-small cell lung cancer patients showed that after including PIWIL1 expression, gender and tumor stage in the multivariable analysis, PIWIL4 failed to remain a significant predictor of mortality (Navarro et al., 2015). On the other hand, three studies on clear cell renal carcinoma, gastric cancer and hepatocellular carcinoma reported no significant association between mortality and PIWIL4 expression. One of these studies showed that neither cytoplasmic nor nuclear nor combined PIWIL4 expression are significantly associated with mortality (Zeng et al., 2017).

We included four studies with 339 participants in the meta-analysis of mortality HR in high compared to low PIWIL4 expression (Figure 8). The pooled HR of mortality in high compared to low PIWIL4 expression was 0.44 (CI 95%: 0.25-0.76, p < 0.05). Minimal heterogeneity was observed in this meta-analysis (I^2^ = 49%, p = 0.12). Sensitivity analysis showed that after removing the study that compared detectable and undetectable rather than high and low PIWIL4 expression, the pooled HR decreased to 0.37 (0.22-0.61, p < 0.05). Publication bias was not assessed in this meta-analysis due to the small number of included studies. Only one study which was performed on patients with non-small cell lung carcinoma investigated the prognostic value of PIWIL4 expression for predicting recurrence and showed no significant association.

**Figure 8.**
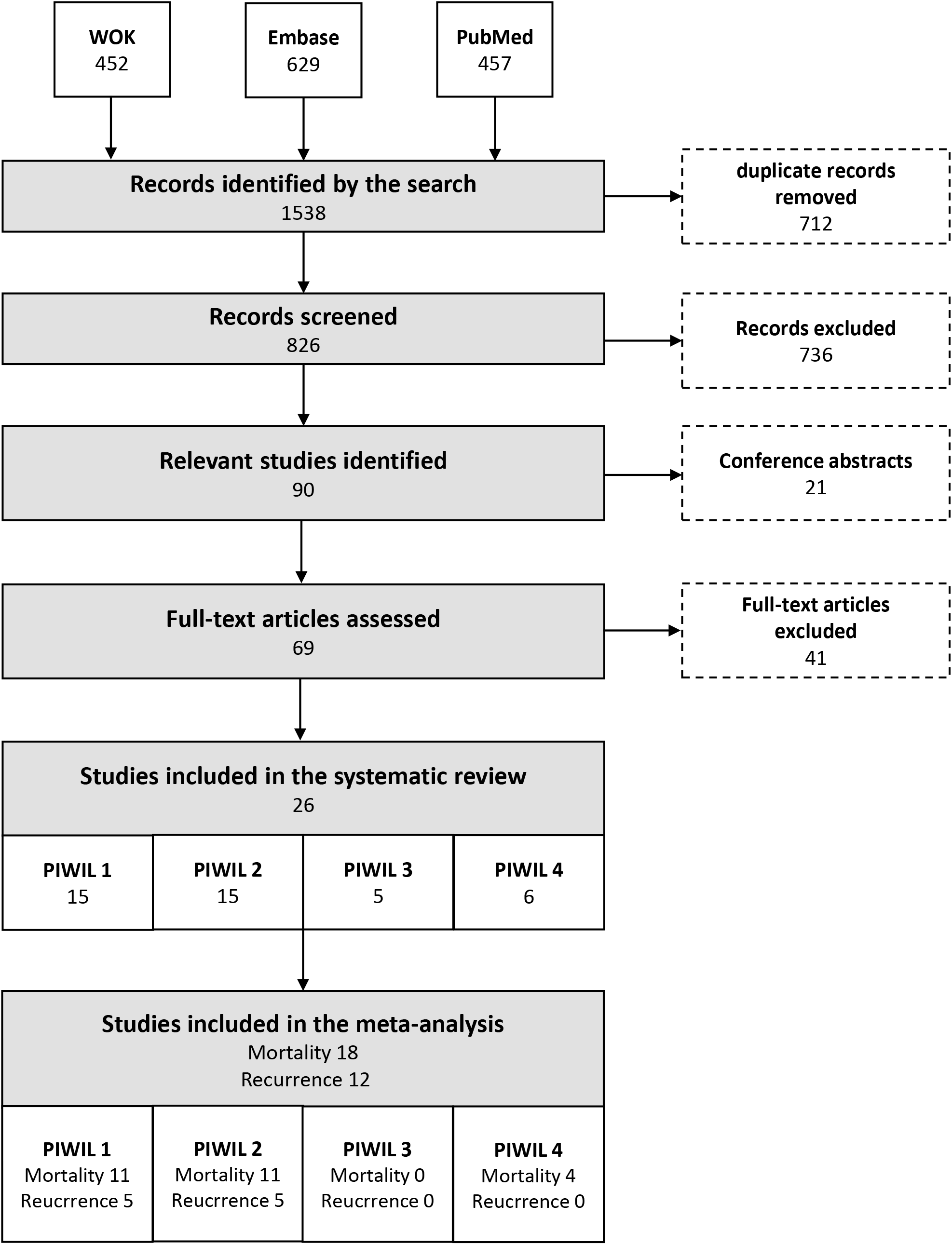
Meta-analysis of HR for mortality in high compared to low expression of PIWIL4 (339 patients). Prognostic value of PIWIL4 for mortality in individual studies are shown in each row. The hazard ratios for individual studies are denoted by squares bounded by a 95% confidence interval bar. The pooled hazard ratio is denoted by a red diamond bounded by a 95% confidence interval bar. Higher HR represents higher risk of mortality in patients with high PIWIL4 expression.

In the TCGA Atlas, PIWIL4 showed highest overall expression levels with many cancer types exhibiting median expression above 1 FPKM. Elevated PIWIL4 expression was noted in stomach and colorectal cancer, as well as in pancreatic cancer. Notably, PIWIL4 expression in testicular cancer was lower than the above 3 cancer types, in contrast to PIWIL1-3 gene expression, where testicular cancer is consistently the highest expressing cancer type (Supplementary Figure 4).

### Combinations of PIWI proteins

The prognostic value of different combinations of PIWI proteins was investigated in two studies. Zeng et al. studied the combination of HILI and PIWIL4 cytoplasmic and nuclear expressions in hepatocellular carcinoma and found no survival difference (Zeng et al., 2017). On the other hand, Greither et al. studied the same combination in female patients with soft tissue sarcoma and showed a better prognosis in patients who have both high HILI and PIWIL4 compared to patients with one gene elevated or both downregulated (Greither et al., 2012). They also studied combination of HILI and PIWIL3 in male patients as well as the combination of HILI, PIWIL3 and PIWIL4 in both genders and showed a lower risk of mortality when all these genes are overexpressed. However, no combination of PIWI proteins was studied in three or more studies, i.e., the minimum required number to perform a meta-analysis; hence, conducting a meta-analysis on different combinations could not be performed.

### Quality of the included studies

Median quality score of the included studies was 55% (range: 35 – 68%) and each study failed to satisfy six quality items on average (range: 4-9). Most of the studies failed to report the inclusion and exclusion criteria, list candidate variables, specify the rationale for their sample size and report the flow of the patients (Table 3).

### Re-analysed survival data from the *Pathology Atlas of The Human Cancer Transcriptome*

We found in some cancer types PIWIL genes to be predictive of patient survival at the formal significance threshold of p value < 0.05 (data not shown). However, the effect on survival for any of the four genes failed to reach the stringent significance threshold, as set by the Atlas creators, of p-value < 0.001 (data not shown).

## DISCUSSION

Here, we showed that increased intratumoral expression of HIWI is linked to higher risk of mortality and recurrence. In contrast, we showed a lower risk of mortality but not recurrence in patients with higher PIWIL4 expression. HILI and PIWIL3 were shown to have no significant association with either mortality or recurrence. In addition, our re-analysis of the TCGA data revealed the different tissues in which the four PIWI proteins are highly expressed in cancer. Therefore, the expression of these proteins can be used for personalized prognostication, which will could be valuable to select the more appropriate treatment for individual patients; so, further studies are needed to establish their role as prognostic marker, notably in tissues marked by high PIWI protein expression levels (such as testis).

Of note, HIWI expression was reported to be an independent prognostic variable when tumor stage, differentiation grade, tumor size, lymph node involvement or PIWIL4 expression were considered in the multivariable models. However, PIWIL4 expression was not significantly associated with survival when it entered the multivariable analysis along with HIWI expression. Furthermore, the combination of PIWI4 with HILI and PIWIL3 was shown to be advantageous in soft tissue sarcomas, while its combination with HILI was not significantly associated with survival in hepatocellular carcinoma.

While the inhibition of HIWI expression decreases the proliferation of gastric cancer cells (although potential effects of *H. pylori*, the major risk factor of gastric cancer (Mentis and Dardiotis, 2019), on PIWI levels have not yet been investigated, to our knowledge (Mentis et al., 2019)), its high expression induces apoptosis in leukemia cell lines (Liu et al., 2006, Sharma et al., 2001). Hence, some included studies hypothesized that there is a U-shaped relationship between PIWI expression and survival. However, our meta-analysis showed no significant difference in mortality between intermediate and high or low HIWI expression. Also, mortality was inversely related to HIWI expression in the case of renal cell carcinoma, unlike other cancers. However, because the number of studies per cancer site is very limited, it is difficult to conclude if this observation is due to that studies included in this meta-analysis are quite divergent, or due to pathophysiological background.

Mechanistically, PIWI proteins bind piRNAs to form a ribonucleoprotein complex that may contribute to tumorigenesis by different mechanisms via silencing transposable elements (Tan et al., 2015). PIWIL2 combination with piR-932 was shown to promote methylation of Latexin, a tumor suppressor gene, in breast cancer cells (Zhang et al., 2013). Furthermore, the inhibition of PIWIL4 and piR-651 complex in gastric cancer cells arrested the cell cycle at the G2/M phase, suggesting a cell proliferative role for this complex (Cheng et al., 2011). Additionally, HIWI can promote indefinite sarcoma cellular proliferation by decreasing cellular differentiation state (Siddiqi et al., 2012). Another study showed that PIWIL2 is involved in DNA repair through relaxation of chromatin (Yin et al., 2011). Mechanistic implications through double-strand DNA breaks production may be also relevant (Thomson and Lin, 2009).

Collectively, previous studies have proposed the involvement of PIWI proteins in several hallmarks of cancer (as proposed by Weinberg and Hanahan (Hanahan and Weinberg, 2000, Hanahan and Weinberg, 2011)) and (reviewed in (Suzuki et al., 2012, Tan et al., 2015)). However, the detailed mechanisms of how PIWI proteins contribute to these hallmarks, and the specific piRNAs involved in these mechanisms are not fully understood (Tan et al., 2015). In addition, the role of PIWI proteins on cancer metabolism, which plays pivotal role in oncogenesis (reviewed in (Mentis and Kararizou, 2010)), appears under-investigated.

The prognostic effect of nuclear and cytoplasmic PIWI expressions was separately investigated in some studies. In patients with esophageal squamous cell carcinoma, cytoplasmic but not nuclear HIWI expression was associated with mortality (He et al., 2009). Similarly, cytoplasmic but not nuclear HILI expression was reported as a predictor of mortality in bladder cancer (Taubert et al., 2015). PIWI proteins may have different roles in nucleus and cytoplasm. For example, while they act as regulators of transposon activity in the nucleus, they regulate RNA stability and translation in the cytoplasm (as discussed in (Taubert et al., 2015)).

In our systematic review, the number of cancer patients with stages I and II cancer approximated the number of patients with stages III and IV cancer. However, a considerable number of studies had not reported the staging. Also, at the individual studies level, this even distribution between early and advanced cases might not be the case, and the results could have been affected by an uneven distribution of patients if their staging was further concerned.

We observed that the prognostic effect of PIWI proteins in different cancer types was highly variable. For example, while the risk of mortality in patients with high compared to low HIWI expression was four-fold in non-small cell lung cancer, it was 69% lower in clear cell renal carcinoma. Even though PIWI proteins are implicated in the very basic mechanisms of cancerogenesis, their effects might be tumor or tissue specific. Besides, their expression in different cancer tissue sites can be substantively different, as our reanalysis of TCGA data revealed. The reasons underlying this variability is unclear and remains to be further investigated but we speculate that perhaps the different piRNAs and the nature of the genes they regulate (i.e., whether they enhance or suppress tumorigenesis) determines the role of PIWI proteins. Furthermore, a combination of different proteins might be at play that influence the role of PIWI proteins in different cancer cells. For example, overexpression of HIWI was shown to have a significant prognostic effect in epithelial ovarian carcinoma only when Let-7a was also overexpressed (Lu et al., 2016).

Nevertheless, our systematic review and meta-analysis had several limitations: (a). The few numbers of included studies for each PIWI protein limited our ability to perform meta-regressions and subgroup analyses; (b). In cases of less than 10 publications, we did not assess the publication bias due to the small number of included studies; (c). We were unable to perform separate analyses for different types of cancers; hence, the analyses should be considered with some caution regarding specific cancers. Indeed, in this meta-analysis, the only cancer site – PIWI protein pair that was investigated in three or more studies, was the prognostic value of HIWI in colorectal cancer recurrence; (d). Proceeding first with the individual patient data meta-analysis, i.e., the gold standard in time-to-event outcomes meta-analyis, was deemed laborious for our study’s capacity; however, aggregated data meta-analyses are also considered of merit (Tudur et al., 2001); (e). Few studies performed multivariable analyses, and the effect sizes included in the meta-analysis were predominantly univariable HRs that we had estimated using Kaplan-Meier plots data. However, our subgroup analyses found that for existing data, there was minimal difference between univariable and multivariable pooled HRs (data not shown). (f). Likewise, the primary meta-analysis on similar survival formats (e.g., OS and DSS) was performed together, because of the limited number of studies; however, subgroup analyses based on type of the outcome for each meta-analysis was performed to explore if the results of meta-analysis for different types of the outcome (e.g. OS and DSS) are significantly different. In most cases, either no significant difference was observed or/and the number of studies per subgroup was less than 3 (data not shown). In every case, though, attention is required, as a recent umbrella review concluded that the “surrogates” may have modest correlation with OS (Haslam et al., 2019); (g). We limited our systematic review to English-language articles and excluded conference abstracts; however, the effect of publication bias in our meta-analysis was negligible. The majority of the included studies were performed in China and Germany, while no studies were performed in America, Africa and Oceania, making the results less representative of the global population. (h). Although not statistically significant, the prognostic role of PIWI proteins expression assessed using RT-PCR or IHC was, in some cases, different; thus, the limitation that different methods to assess the presence of PIWI proteins (e.g., immuno-cytochemistry, real-time PCR, PCR, ELISA) may have different accuracy and precision (e.g., sensitivity and specificity) levels should be acknowledged. Similarly, it is well known that not all antibodies are the same with respect to selectivity and specificity; this can be a factor contributing to the different results observed among the studies. However, it is difficult to clearly appreciate the analytical validity of the antibodies used in the included studies that measured PIWI expression by IHC. In parallel, the difference in mortality risk of high compared to low PIWI expression was generally underestimated when its mRNA expression was measured compared to its protein expression. PIWI proteins translated from their mRNAs might undergo different post-translational modification or alternative splicing which can influence the IHC results. For example, there are five different protein isoforms for HILI, including PL2L80, PL2L60, PL2L50, PL2L42 and PL2L40 (Ye et al., 2010). Furthermore, different scoring methods in IHC as well as different cut-off values for mRNA levels used to classify the patients to high and low PIWI expression, might also lead to under- or over-estimation of its expression and influence the results. Hence, the potential underlying difference between mRNA and protein expression levels *per se* and of the drawbacks in their measurement, which are not always parallel, should be acknowledged while interpreting the study’s results. Therefore, the subgroup analysis for each PIWI protein based on the method of PIWI expression measurement is essential. (i). Last, the outcomes deduced from the meta-analyses can be very sensitive, besides the sample size effects, to confounding variables used in each study and other factors and such as race, geographical location of the included study, duration of a given study; hence, caution in interpretation and conducting larger validation studies are both needed before PIWI proteins reach clinical prime-time as biomarkers in our precision medicine era (Mentis et al., 2018).

## CONCLUSIONS

To our knowledge, this systematic review and meta-analysis is the first one attempted on the prognostic role of PIWI proteins on cancer. We reported an overall worse prognosis in patients with higher HIWI and lower PIWIL4 expression. Additional studies are needed to investigate and validate the prognostic effect of PIWI proteins in different cancers. Crucially, future studies should be performed and make their data publicly available, so that large analyses investigating a combination of different markers, such as the Pathology Atlas (Uhlen et al., 2017) and Kaplan-Meir Plotter (Lánczky et al., 2016), are enriched. Furthermore, because of the PIWI proteins restricted expression in the germ cells and tumors, these proteins can be theorized as good targets for better anti-neoplastic drugs with less side effects (Tan et al., 2015), notably in cancer patients of non-reproductive age, as part of precision medicine approaches.

## Tables Legends

**Table 1. Characteristics of the studies included in the meta-analysis.**

**Table 2. Descriptive Statistics of Eligible Studies**

**Table 3. Quality of the included studies assessed using REMARK checklist.** Y: fully satisfied, P: partially satisfied, N: not satisfied, NA: not applicable, U: unclear

**Table S1. Search strategies used for different databases**

---

**Supplementary Figure 1.** Expression of PIWI-1 protein across various cancer types, as per the Human Protein Pathology Atlas

**Supplementary Figure 2.** Expression of PIWI-1 protein across various cancer types, as per the Human Protein Pathology Atlas

**Supplementary Figure 3.** Expression of PIWI-3 protein across various cancer types, as per the Human Protein Pathology Atlas

**Supplementary Figure 4.** Expression of PIWI-4 protein across various cancer types, as per the Human Protein Pathology Atlas

