## Supplementary material for "PIWI proteins as prognostic markers in cancer: a systematic review and meta-analysis": Table 1

| **Table 1. Characteristics of the studies included in the meta-analysis.** | | | | | | | | | | |
| --- | --- | --- | --- | --- | --- | --- | --- | --- | --- | --- |
| **First Author** | **Country** | **Number of participants** | **Cancer type** | **Mean or median follow-up duration (months)** | **Mortality outcome type** | **Recurrence outcome type** | **Diagnostic method** | **Type of PIWIL** | **Mortality results** | **Recurrence results** |
| (Al-Janabi et al., 2014) | Germany | 73 | clear cell renal cell carcinoma | 24 | OS, DSS | - | RT-PCR | 1 | Statistical method not specified: not significant |  |
|  |  |  |  |  |  |  |  | 2 | Statistical method not specified: not significant |  |
|  |  |  |  |  |  |  |  | 3 | Statistical method not specified: not significant |  |
|  |  |  |  |  |  |  |  | 4 | Statistical method not specified: not significant |  |
| (Cao et al., 2016) | China | 187 | breast cancer | 100 | NR | - | RT-PCR | 1 | log-rank: p < 0.05 HR uHRc: 2.14 (1.27-3.63) |  |
|  |  |  |  |  |  |  |  | 2 | log-rank: not significant uHRc: 1.05 (0.61-1.79) |  |
| (Chen et al., 2015) | China | 41 | hilar cholangiocarcinoma | 26.2 | OS | DFS | IHC | 2 | log-rank: p < 0.05 mHR: 3.95 (1.10-10.98) | log-rank: p < 0.05 mHR: 4.04 (1.09-11.36) |
| (Gambichler et al., 2017) | Germany | 183 | malignant melanoma | NR | NR | NR | IHC | 3 | Statistical method not specified: not significant | Statistical method not specified: not significant |
| (Greither et al., 2012) | Germany | 125 | soft tissue sarcoma | 32 | DSS | NR | RT-PCR | 2 | mRR: 0.53 (0.30-0.94) |  |
|  |  |  |  |  |  |  |  | 3 | mRR: 0.75 (0.43-1.33) |  |
|  |  |  |  |  |  |  |  | 4 | mRR: 0.54 (0.31-0.97) |  |
|  |  |  |  |  |  |  |  | 2 & 4 | log-rank (females): p < 0.05 mRR: 0.38 (0.18-0.79) |  |
|  |  |  |  |  |  |  |  | 2 & 3 | log-rank (males): p < 0.05 mRR: 0.47 (0.22-0.99) |  |
|  |  |  |  |  |  |  |  | 2 & 3 & 4 | mRR: 0.24 (0.09-0.62) |  |
| (Grochola et al., 2008) | Germany | 78 | ductal adenocarcinoma of the pancreas | 15.99 mean (range 1–61) months | DSS | - | IHC | 1 | Statistical method not specified: not significant |  |
|  |  | 56 |  |  |  |  | RT-PCR |  | mRR (high vs low): not significant mRR (detectable vs non-detectable): 1.40 (CI NR) mRR (intermediate vs high or low in males): 2.78 (CI NR) mRR (intermediate vs high or low in females): not significant |  |
| (He et al., 2009) | China | 153 | esophageal squamous cell carcinoma | 124 months (range 118-155 months) | DSS | - | IHC | 1 | log-rank (high and moderate vs low): p < 0.05 log-rank (high vs moderate): p = 0.60 uHRc (intermediate vs high or low): 1.81 (1.16-2.80) mRR (cytoplasm): 2.25 (1.37-3.67) mRR (nucleus): 1.23 (0.76-1.99) |  |
| (Iliev et al., 2016) | Czech Republic | 57 | clear cell renal cell carcinoma | NR | OS | - | RT-PCR | 1 | log-rank: p < 0.05 uHRc: 0.31 (0.15-0.63) |  |
|  |  |  |  |  |  |  |  | 2 | log-rank: p < 0.05 uHRc: 0.14 (0.07-0.27) |  |
|  |  |  |  |  |  |  |  | 3 | log-rank: p = 0.85 |  |
|  |  |  |  |  |  |  |  | 4 | log-rank: p < 0.05 uHRc: 0.28 (0.14-0.57) |  |
| (Li et al., 2012) | China | 185 [203 total] | colon cancer | 71.6 mean (range: 63.7-84.7) months | OS | MFS | IHC | 2 | log-rank: p < 0.05 uHR: 3.31 (1.59–6.90) mHR: 4.33 (2.05–9.13) | log-rank: p < 0.05 uHR: 2.63 (1.17–5.81) mHR: 4.03 (1.74–9.33) |
| (Li et al., 2017) | China | 97 | glioma | NR | OS | - | IHC | 2 | log-rank: p < 0.05 uHR: 4.78 (2.32-9.85) |  |
| (Litwin et al., 2018) | Poland | 101 | breast cancer | NR | NR | - | IHC | 1 | log-rank: p = 0.41 uHRc: 2.36 (0.66-8.49) |  |
|  |  |  |  |  |  |  |  | 2 | log-rank: p = 0.44 uHRc: 2.27 (0.75-6.82) |  |
| (Litwin et al., 2018) | China | 225 | colon adenocarcinoma | 45.0 + 28.4 (range 1-120) months | OS | DFS | IHC | 1 | uHR: 1.1 (0.6–2.2) |  |
| (Lu et al., 2016) | Italy | 211 | epithelial ovarian cancer | 31 (0.6-114) months | OS | DFS | RT-PCR | 1 | uHR (intermediate vs high or low): 1.63 (1.06–2.52) mHR (intermediate vs high or low): 1.89 (1.29–2.98) | uHR (intermediate vs high or low): 1.33 (0.863–2.05) mHR (intermediate vs high or low): 1.38 (0.88–2.16) |
| (Lu et al., 2016) | Spain | 71 | non-small cell lung cancer | NR | OS | RFS | IHC | 1 | log-rank: p < 0.05 uHRc: 3.73 (1.43-9.73) | log-rank: p < 0.05 mHR: 2.89 (1.13–7.39) |
|  |  |  |  |  |  |  | RT-PCR | 4 | log-rank: p < 0.05 uHRc: 0.24 (0.08-0.69) | log-rank: p < 0.05 uHRc: 0.53 (0.20-1.41) mHR: NR, p = 0.09 |
| (Oh et al., 2012) | South Korea | 60 | colorectal carcinoma | 28 months (range, 2 to 72 months) | OS | - | IHC | 2 | log-rank: p = 0.05 | uHR: NR, p = 0.05 mHR: NR, p = 0.60 |
| (Pouyanfar et al., 2016) | Iran | 35 | prostate cancer | 8 years [no sd or range] | OS | - | RT-PCR | 2 | Statistical method not specified: p < 0.05 |  |
| (Qu et al., 2015) | China | 126 | non-small cell lung cancer | 100 months [no sd or range] | OS | DFS | RT-PCR | 2 | log-rank: p < 0.05 uHRc: 1.96 (1.15-3.34) | log-rank: p < 0.05 uHRc: 1.55 (0.91-2.63) |
| (Sun et al., 2011) | China | 66 | glioma | NR | DSS | - | IHC | 1 | log-rank: p < 0.05 uHRc: 2.86 (1.59-5.14) |  |
| (Sun et al., 2017) | China | 110 | colorectal carcinoma | NR | OS | DFS | IHC | 1 | log-rank: p < 0.05 mHR: 2.80 (1.37–5.72) | log-rank: p < 0.05 mHR: 3.23 (1.56–6.66) |
| (Taubert et al., 2007) | Germany | 65 | soft tissue sarcoma | 65 mean (SD NR) | NR | - | RT-PCR | 1 | log-rank (high vs intermediate): p < 0.05 log-rank (low vs intermediate): p = 0.17 uHRc (intermediate vs high or low): 0.39 (0.17-0.88) |  |
| (Taubert et al., 2015) | Germany | 202 | bladder cancer | NR | DSS | PFS | IHC | 2 | log-rank: p < 0.05 uHRc: 0.24 (0.14-0.41) | log-rank: p = 0.07 uHRc: 0.38 (0.23-0.63) |
| (Wang et al., 2012) | China | 182 | gastric cancer | 35 median months | OS | - | IHC | 1 | log-rank: p < 0.05 mHR: 2.66 (1.10-6.45) |  |
|  |  |  |  |  |  |  |  | 2 | log-rank: p < 0.05 mHR: 0.83 (0.45-1.51) |  |
|  |  |  |  |  |  |  |  | 3 | log-rank: p = 0.36 |  |
|  |  |  |  |  |  |  |  | 4 | log-rank: p = 0.55 |  |
| (Zeng et al., 2017) | China | 86 | hepatocellular carcinoma | 4-7 years died: 14 (1-95) months survived: 57 (46-80) months | NR | - | IHC | 2 | log-rank: p = 0.52 uHRc: 0.99 (0.56-1.74) |  |
|  |  |  |  |  |  |  |  | 4 | log-rank: p = 0.29 uHRc: 0.92 (0.37-2.24) |  |
|  |  |  |  |  |  |  |  | 2+4 | log-rank: p = 0.21 |  |
| (Zeng et al., 2011) | China | 270 | colorectal carcinoma | 33 (3-76) months | OS | DFS | IHC | 1 | log-rank: p = 0.12 mHR: 1.67 (1.13–2.48) | log-rank: p = 0.15 uHRc: 1.32 (0.91-1.92) |
| (Zhang et al., 2013) | China | 1086 | breast cancer | NR | DSS | MFS | IHC | 2 |  | mHR: 5.64 (3.57-9.58) |
| (Zhao et al., 2012) | China | 168 | hepatocellular carcinoma | 31.6 median | OS | RFS | IHC | 1 | log-rank: p < 0.05 mHR: 1.88 (1.19-2.95) | log-rank: p < 0.05 mHR: 1.72 (1.03-2.85) |
