## Supplementary material for "PIWI proteins as prognostic markers in cancer: a systematic review and meta-analysis": Table 2

**Table 2. Descriptive Statistics of Eligible Studies**

| ***Characteristic*** | ***Subgroup*** | ***Frequency (%)*** |
| --- | --- | --- |
| *PIWI protein* | PIWIL1 (HIWI) | 15 (58%) |
|  | PIWIL2 (HILI) | 15 (58%) |
|  | PIWIL3 | 5 (19%) |
|  | PIWIL4 | 6 (23%) |
| *Country of study* | China | 14 (54%) |
|  | Germany | 6 (23%) |
|  | Czech Republic | 1 (4%) |
|  | Iran | 1 (4%) |
|  | Italy | 1 (4%) |
|  | Poland | 1 (4%) |
|  | South Korea | 1 (4%) |
|  | Spain | 1 (4%) |
| *Cohort start decade* | 1980s | 1 (4%) |
|  | 1990s | 3 (12%) |
|  | 2000s | 12 (46%) |
|  | 2010s | 1 (4%) |
|  | NR | 9 (35%) |
| *Cancer site* | colon and rectum | 5 (19%) |
|  | breast | 3 (12%) |
|  | kidney | 2 (8%) |
|  | liver | 2 (8%) |
|  | lung | 2 (8%) |
|  | brain | 2 (8%) |
|  | soft tissue | 2 (8%) |
|  | bladder | 1 (4%) |
|  | esophagus | 1 (4%) |
|  | gallbladder | 1 (4%) |
|  | ovaries | 1 (4%) |
|  | pancreas | 1 (4%) |
|  | prostate | 1 (4%) |
|  | stomach | 1 (4%) |
| *Outcome studied* | OS | 15 (58%) |
|  | DSS | 8 (30%) |
|  | unspecified mortality | 5 (19%) |
|  | DFS, PFS | 6 (23%) |
|  | MFS | 2 (8%) |
|  | RFS | 2 (8%) |
|  | unspecified recurrence | 2 (8%) |
| *Quantification method* | IHC | 18 (69%) |
|  | RT-qPCR | 8 (30%) |
| *Average duration of follow-up* | < 2 years | 1 (4%) |
|  | 2 to 4 years | 9 (35%) |
|  | 4 to 6 years | 3 (12%) |
|  | > 6 years | 4 (15%) |
|  | NR | 9 (35%) |
