## Supplementary material for "PIWI proteins as prognostic markers in cancer: a systematic review and meta-analysis": Table 3

| **Table 3. Quality of the included studies assessed using REMARK checklist. Y: fully satisfied, P: partially satisfied, N: not satisfied, NA: not applicable, U: unclear** | | | | | | | | | | | | | | | | | | | | | | | | | | | |
| --- | --- | --- | --- | --- | --- | --- | --- | --- | --- | --- | --- | --- | --- | --- | --- | --- | --- | --- | --- | --- | --- | --- | --- | --- | --- | --- | --- |
| **#** | Item Described | Al-Janabi, O. (2014) | Cao, J. (2016) | Chen, Y. J. (2015) | Gambichler, T. (2017) | Greither, T. (2012) | Grochola, L. F. (2008) | He, W. (2009) | Iliev, R. (2016) | Li, D. (2012) | Li, J. (2017) | Litwin, M. (2018) | Liu, C. (2012) | Lu, L. (2016) | Navarro, A. (2015) | Oh, S. J. (2012) | Pouyanfar, N. (2016) | Qu, X. (2015) | Sun, G. (2011) | Sun, R. (2017) | Taubert, H. (2007) | Taubert, H. (2015) | Wang, Y. (2012) | Zeng, G. (2017) | Zeng, Y. (2011) | Zhang, H. (2013) | Zhao, Y. M. (2012) |
| **1** | Study objectives | Y | Y | Y | Y | Y | Y | Y | Y | Y | Y | Y | Y | Y | Y | Y | Y | Y | Y | Y | Y | Y | Y | Y | Y | Y | Y |
| **2** | Patients characteristics | P | N | N | N | N | P | N | N | P | N | N | N | P | N | P | Y | P | P | N | N | P | N | N | N | Y | N |
| **3** | Treatment details | N | N | P | N | N | P | P | P | P | P | P | P | P | Y | P | Y | N | P | P | N | P | N | P | Y | P | N |
| **4** | Biological material and method of preservation | Y | N | N | P | P | Y | Y | N | P | Y | P | P | N | Y | P | P | N | P | P | P | P | P | Y | P | Y | P |
| **5** | Assay methods | Y | Y | Y | Y | Y | Y | Y | Y | Y | Y | Y | Y | Y | Y | Y | Y | Y | Y | Y | P | Y | Y | Y | Y | P | Y |
| **6** | Study design (prospective or retrospective) | P | P | P | N | P | P | Y | N | P | N | N | P | P | P | Y | P | P | N | N | P | N | P | P | P | N | P |
| **7** | Endpoints definition | P | P | P | P | P | P | P | P | P | P | N | P | P | P | Y | Y | P | P | P | N | P | Y | Y | Y | P | Y |
| **8** | List candidate variables | N | N | N | N | N | Y | Y | N | N | N | N | N | N | N | Y | N | N | N | N | N | Y | N | N | N | N | N |
| **9** | Rationale for sample size | N | N | N | N | N | N | N | N | N | N | N | N | N | N | N | N | N | N | N | N | N | N | N | N | N | N |
| **10** | Statistical analyses | P | Y | Y | Y | Y | Y | Y | Y | Y | Y | Y | Y | Y | Y | Y | Y | Y | Y | Y | N | Y | Y | Y | Y | Y | Y |
| **11** | Continuous vs binary | Y | P | P | Y | Y | Y | Y | N | Y | Y | Y | Y | Y | Y | Y | N | Y | Y | Y | P | Y | Y | Y | Y | Y | Y |
| **12** | Flow of the patients | N | N | N | N | N | N | N | N | N | N | N | N | N | N | N | N | N | N | N | N | N | N | N | N | N | Y |
| **13** | Demographics | P | Y | Y | Y | Y | Y | Y | Y | Y | Y | Y | Y | Y | Y | Y | Y | Y | Y | Y | P | Y | Y | Y | N | Y | Y |
| **14** | Marker relation to standard prognostic variables | Y | Y | Y | P | Y | P | Y | Y | Y | Y | Y | P | Y | N | Y | Y | Y | N | Y | Y | P | Y | N | N | Y | Y |
| **15** | Univariable analysis | N | P | N | P | P | N | P | P | Y | Y | P | Y | Y | P | P | P | P | P | P | P | Y | P | P | P | N | P |
| **16** | Multivariable analysis | N | N | Y | U | Y | P | P | N | Y | N | N | N | Y | Y | P | N | N | N | Y | Y | Y | Y | N | Y | Y | Y |
| **17** | Variables included in multivariable analysis regardless of univariable p-value | NA | NA | U | U | U | U | Y | NA | U | NA | NA | N | U | N | N | NA | NA | NA | NA | U | Y | U | NA | NA | NA | N |
| **18** | Checking assumptions and sensitivity analyses | N | N | N | N | N | N | N | N | N | N | N | N | N | N | N | N | N | N | N | N | N | N | N | N | N | N |
| **19** | Discussions | Y | P | N | Y | Y | Y | Y | P | Y | Y | Y | Y | Y | Y | Y | P | Y | P | Y | P | Y | Y | P | Y | Y | Y |
| **20** | Implications for research and clinic | P | N | N | Y | Y | Y | P | P | Y | P | P | Y | P | P | P | P | P | Y | Y | P | P | Y | P | Y | Y | Y |
|  | Score | 47% | 39% | 40% | 43% | 55% | 60% | 68% | 39% | 63% | 55% | 47% | 53% | 58% | 55% | 65% | 55% | 50% | 47% | 58% | 35% | 65% | 58% | 50% | 55% | 61% | 63% |
