## Supplemental Table 1 for "PIWI proteins as prognostic markers in cancer: a systematic review and meta-analysis"

| **Database** | **Search strategy** |
| --- | --- |
| PubMed | (PIWI[tiab] OR PIWIL*[tiab] OR HIWI[tiab] OR HILI[tiab]) AND (cancer*[tiab] OR carcinoma*[tiab] OR tumor[tiab] OR tumour[tiab] OR tumors[tiab] OR tumours[tiab] OR neoplas*[tiab] OR malignan*[tiab] OR metasta*[tiab] OR leukemia[tiab] OR leukaemia[tiab] OR lymphoma[tiab]) |
| Embase | (piwi:ti,ab OR 'piwil*':ti,ab OR hiwi:ti,ab OR hili:ti,ab) AND ('cancer*':ti,ab OR 'carcinoma*':ti,ab OR 'tumor':ti,ab OR 'tumour':ti,ab OR 'tumors':ti,ab OR 'tumours':ti,ab OR 'neoplas*':ti,ab OR 'malignan*':ti,ab OR 'metasta*':ti,ab OR 'leukemia*':ti,ab OR 'leukaemia*':ti,ab OR 'lymphoma*':ti,ab) |
| Web of Knowledge* | (PIWI OR PIWIL* OR HIWI OR HILI) AND (cancer* OR carcinoma* OR tumor OR tumour OR tumors OR tumours OR neoplas* OR malignan* OR metasta* OR leukemia OR leukaemia OR lymphoma) |

**Table S1. Search strategies used for different databases**

* in topic
