## Supplementary figures and images for "PIWI proteins as prognostic markers in cancer: a systematic review and meta-analysis"

### Supplementary Figure 1.png

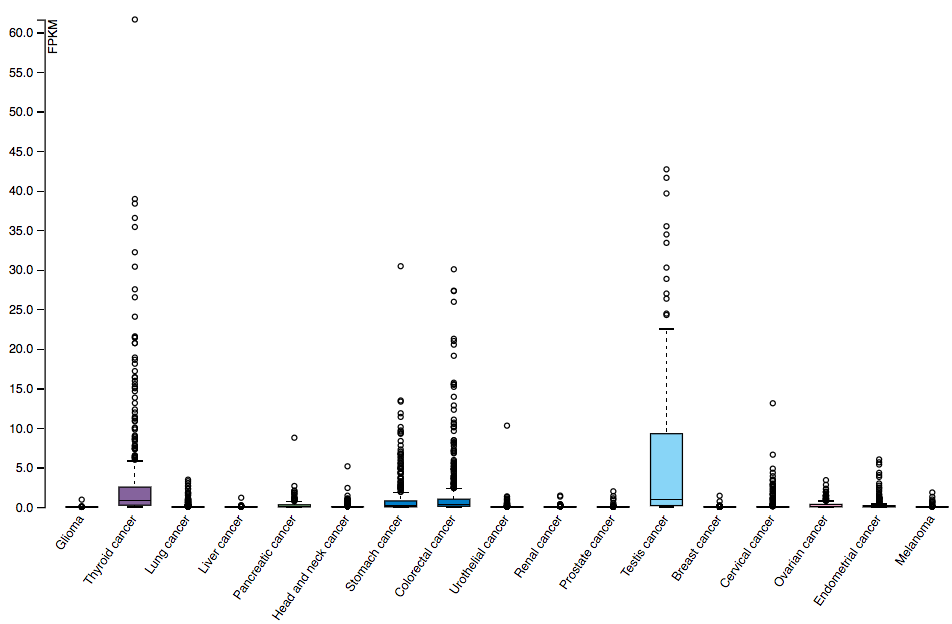

### Supplementary Figure 2.png

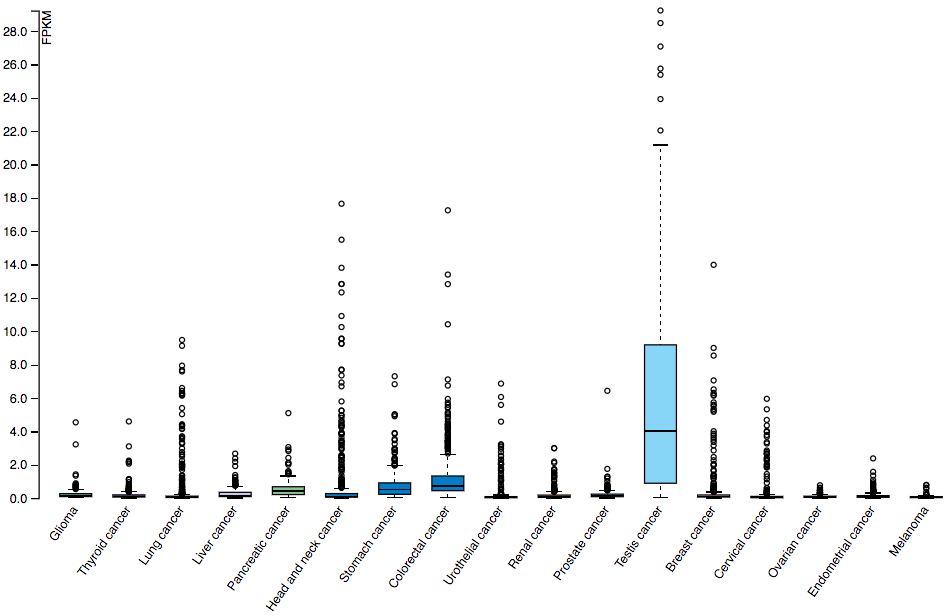

### Supplementary Figure 3.png

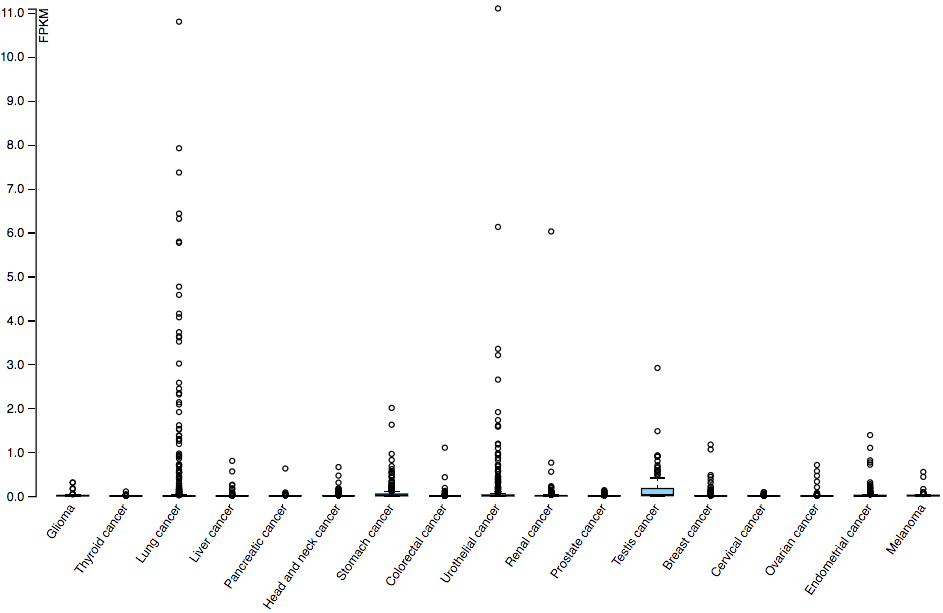

### Supplementary Figure 4.png

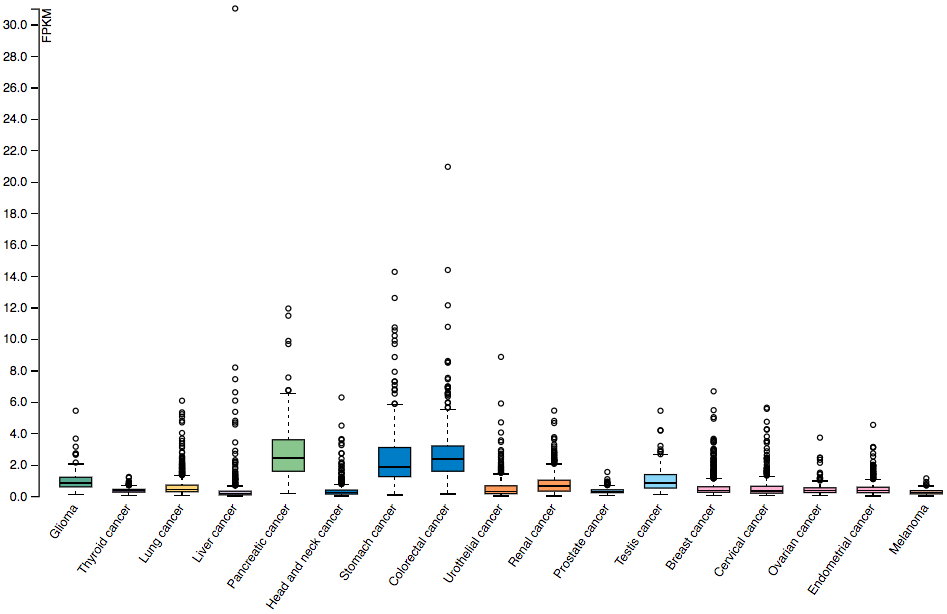
